# Quantitative trait loci mapping of gene expression and chromatin accessibility in primary fibroblast reveals shared allelic effects between Latin American and European ancestries

**DOI:** 10.1101/2025.06.09.658613

**Authors:** Toni Boltz, Merel Bot, Sandra Lapinska, Tommer Schwarz, Kangcheng Hou, Kristina M. Garske, Malika K. Freund, Carrie E. Bearden, Gabriel Macaya, Carlos Lopez-Jaramillo, Nelson Freimer, Marco P. Boks, Rene S. Kahn, Bogdan Pasaniuc, Roel A. Ophoff

**Affiliations:** Department of Human Genetics, David Geffen School of Medicine, UCLA; Center for Neurobehavioral Genetics, Semel Institute for Neuroscience and Human Behavior, UCLA; Graduate Group in Computational Biology, University of Pennsylvania; Department of Bioinformatics, David Geffen School of Medicine, UCLA; Department of Psychology, UCLA; Center for Research in Cellular and Molecular Biology, Universidad de Costa Rica, San José, Costa Rica; Department of Psychiatry, University of Antioquia, Medellín, Antioquia, Colombia; Department of Psychiatry, Amsterdam UMC, Amsterdam, The Netherlands; Dimence Institute for Specialized Mental Health Care, Dimence Group, Deventer, The Netherlands; Department of Psychiatry, Brain Center University Medical Center Utrecht, University Utrecht, Utrecht, the Netherlands; Department of Psychiatry, Icahn School of Medicine at Mount Sinai, New York, NY, USA; Department of Genetics, University of Pennsylvania

## Abstract

Quantitative Trait Locus (QTL) analysis of molecular data has identified genetic variants associated with traits such as gene expression, and colocalization of these functional QTL with GWAS risk loci has offered insights into the genetic basis of human disease. We employed gene expression (RNA-seq) and chromatin accessibility (ATAC-seq) obtained from human primary fibroblasts to investigate quantitative trait loci (QTLs) in cohorts ascertained for bipolar disorder of European (n=150) and Latin American (n=96) ancestries. Leveraging data from three countries of origin (The Netherlands, Colombia, Costa Rica) within our cohort, we characterized differences among individuals at the SNP, gene, and accessible-chromatin levels to compute ancestry-specific expression (e)QTLs and chromatin-accessibility (ca)QTLs. Across ancestries, we observed R² ≥ 0.93 for eQTL effect sizes and R² ≥ 0.95 for caQTLs, indicating a high degree of concordance. Integrating chromatin data with expression and genotype information enabled precise fine-mapping of eQTLs, yielding 203 high-confidence (posterior probability > 90 %) regulatory pathways. In downstream analyses, transcriptome-wide (TWAS) and chromatin-wide (CWAS) association studies with brain- and skin-related GWAS identified 36 TWAS-significant genes and 77 CWAS-significant open chromatin regions. These findings underscore the shared genetic regulatory mechanisms across European and Latin American ancestries, while demonstrating that ancestry-specific reference panels enhance the accuracy of TWAS and CWAS in diverse populations.

## INTRODUCTION

In recent years, genome-wide association studies (GWAS) have reached sample sizes in the millions of individuals, yet nearly 80% of individuals included in these studies are of European descent.^1^ Similarly, large-scale omics resources, such as GTEx, are overwhelmingly comprised of European-ancestry donors.^2^ This skew limits our ability to discover genetic signals that are robust and generalizable across all populations, as variant frequencies and linkage disequilibrium (LD) patterns differ by ancestry,^3,4,5^ emphasizing a need for genetic and genomic datasets from diverse populations. Incorporating data from diverse ancestries allows for the identification of both shared and population-specific genetic contributors to various traits, leading to a more comprehensive understanding of the underlying molecular mechanisms.^6^

Furthermore, GWAS have successfully discovered loci associated with the risk of developing various complex diseases, including 298 independent loci currently reported for bipolar disorder.^7^ However, the majority of these loci are in non-coding regions of the genome and as such, the causal mechanism between the genetic variation and disease risk often is unclear. Quantitative trait loci (QTL) studies^8^ have identified non-coding SNPs that impact both gene expression and complex phenotypes, revealing mechanistic insights into disease architecture.^9–11^ While fibroblasts are not the preferred cell type for studying psychiatric disease susceptibility, previous QTL studies have shown that cis-genetic effects are generally shared across cell types, with the caveat that brain tissue types are more highly correlated with each other than non-brain tissue types.^12,13^ Relatedly, previous studies have shown that cis-effects on chromatin accessibility tend to be less context-dependent than on gene expression,^14^ thus we expect better power to detect associations despite the tissue type. Given the ease of accessibility of a skin sample relative to a brain tissue sample, fibroblasts provide a unique opportunity to study biological samples from cohorts of hundreds of individuals. We consider fibroblasts to be a better tissue for studying neuropsychiatric traits than blood given the closer developmental origin (fibroblasts and neurons from the ectoderm, blood from the mesoderm) and greater transcriptomic stability of fibroblasts, compared to various blood cell types that require rapid response to immune or environmental perturbations.

In this study, we perform a QTL mapping study in fibroblasts of a multi-ancestry sample with multi-omics data. Specifically, we investigate human primary fibroblasts derived from skin biopsies, originating from participants from The Netherlands (n=150), Colombia (n=50), and Costa Rica (n=46), with SNP-genotypes, gene expression via RNA-seq, and chromatin accessibility via ATAC-seq^15^ (Assay for Transposon-Accessible Chromatin sequencing) data measured for all individuals. We use this uniquely large and diverse functional genomics dataset to evaluate effects of quantitative traits and their association with GWAS risk loci. We found that eQTL and caQTL effect sizes in fibroblasts are highly concordant across European and Latin American ancestries, enabling fine-mapping of high-confidence regulatory pathways. Downstream TWAS and CWAS identified 36 genes and 77 open-chromatin regions associated with brain- and skin-related GWAS loci, underscoring shared genetic regulation and the value of ancestry-specific reference panels.

## RESULTS

### Population stratification apparent in SNP-genotypes but not in gene expression or chromatin accessibility

Given the three countries of origin within our cohort, we initially characterized the differences amongst these individuals at the SNP, gene, and accessible-chromatin levels, which were used to compute ancestry-specific expression (e)QTLs and chromatin-accessibility (ca)QTLs. Principal component analysis (PCA) of the imputed genotypes depicted clear and significantly different ancestry-specific clusters (PC1 P_ANOVA_ = 5.6e-16, PC2 P_ANOVA_ = 5.7e-14 after correcting for batch year) (Figure 1A) (Table S1), as expected.^16^ See Figure S1 for the PCA on genotypes overlaid with 1000 Genomes reference populations. However, principal component analysis of the gene expression and peak matrices revealed stronger correlations with batch year than with ancestry (Table S1) (Figures 1B and 1C). This suggests that while population stratification is clearly detectable in the SNP-genotype data, it is not as impacted at the transcriptomic or accessible chromatin levels, which are more strongly impacted by batch-related effects. Figure S2 provides the PCA plot for the expression data after correcting for year, ancestry, and other technical factors as described in the **Methods**; similarly, Figure S3 provides the plot of the chromatin data after correction. The residual values after correcting for all technical and batch related covariates for gene expression and chromatin peaks are used in the downstream analyses presented here.

**Figure 1.**
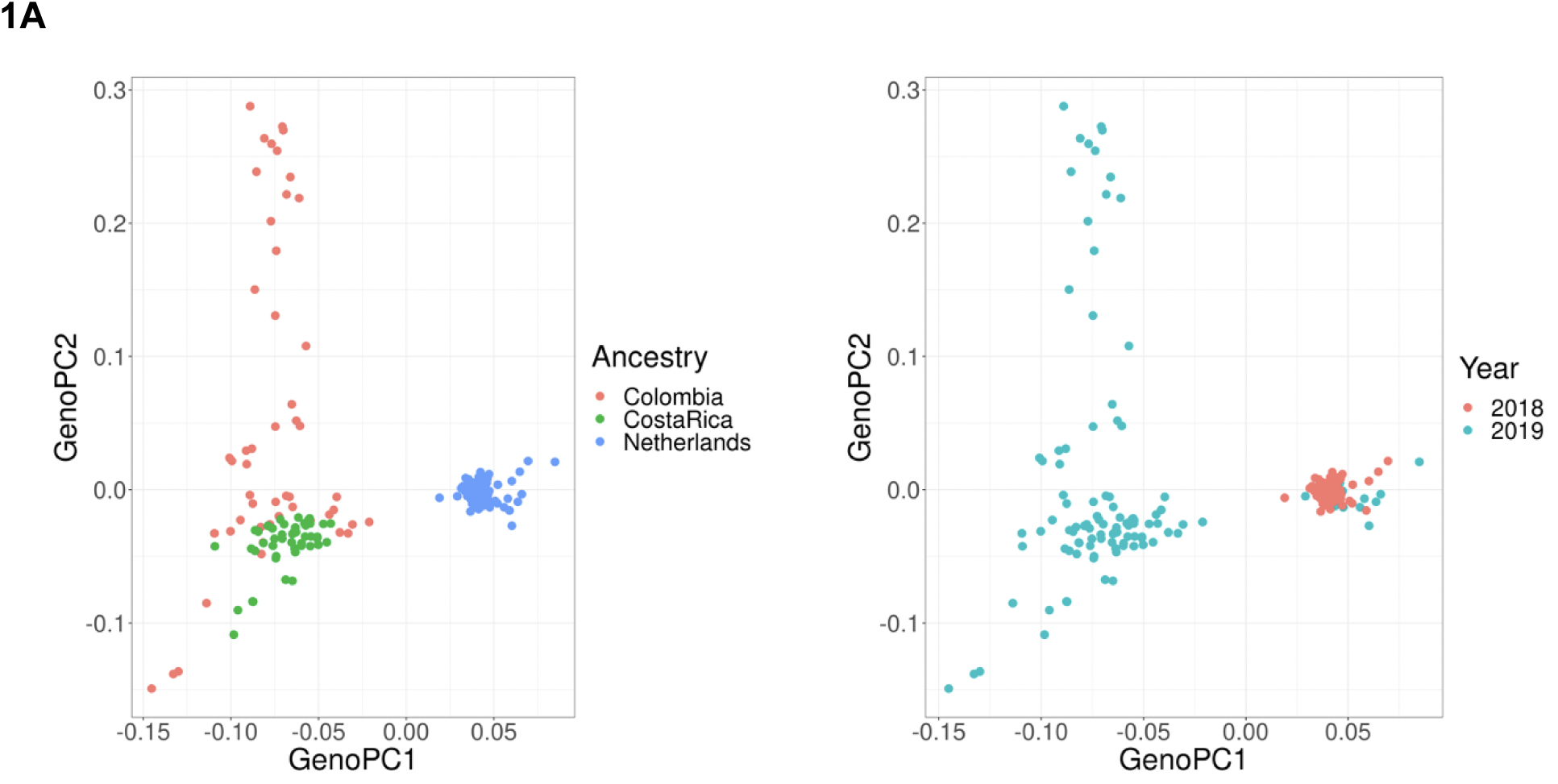
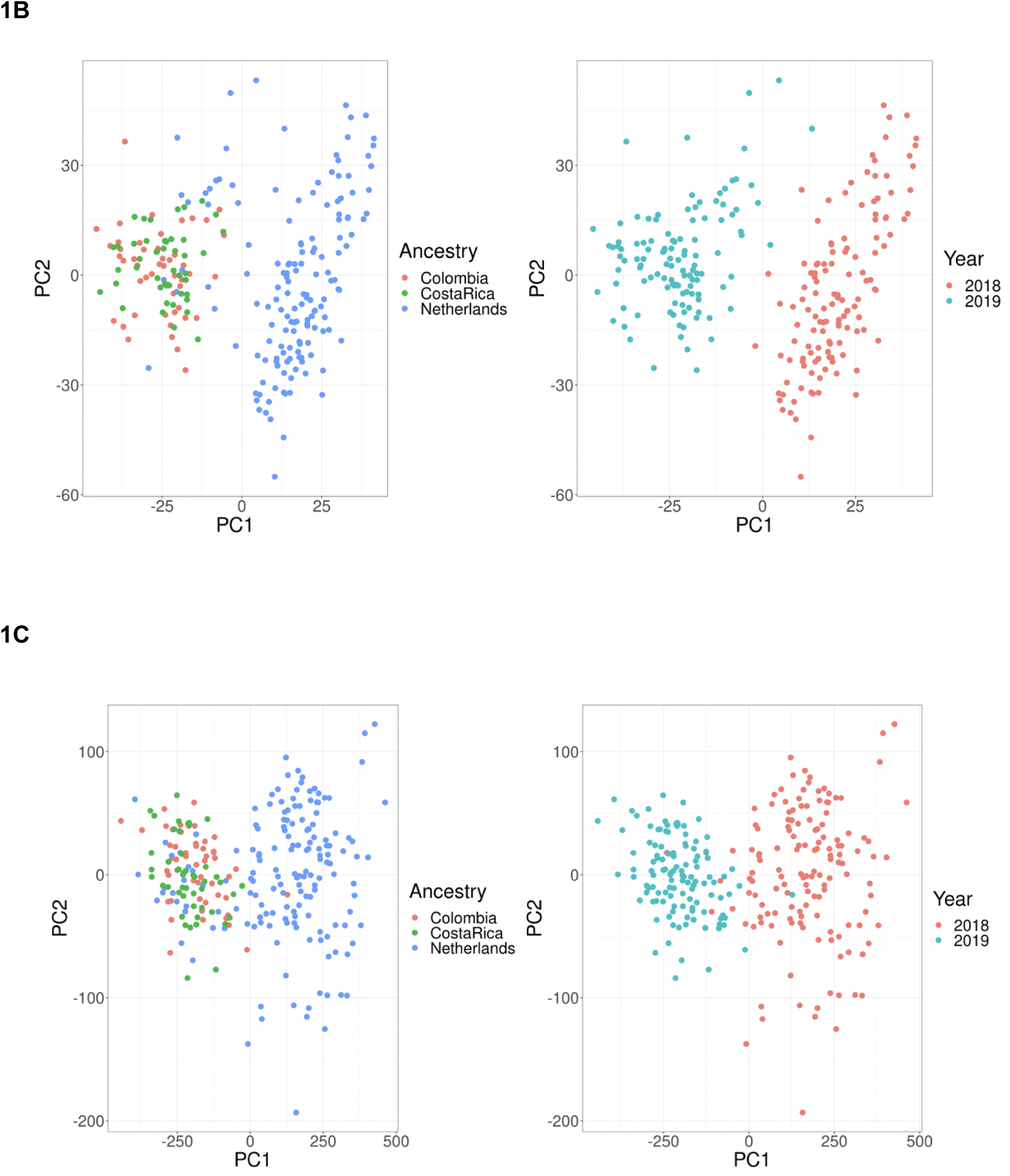
Principal component analysis of genotype, expression, and chromatin datasets. **(A)** PCA on genotypes, each dot represents an individual. The left plot is colored by country of origin and the right plot is colored by sequencing batch year. PC1 explains 4.9% of the variance while PC2 explains 2.8%. **(B)** PCA on gene expression. PC1 explains 10.8% of the variance while PC2 explains 6.8%. Regarding association to ancestry group, we found that PC1 P_ANOVA_ = 0.0027 and PC2 P_ANOVA_ = 0.113 after correcting for batch year. **(C)** PCA on chromatin peaks. PC1 explains 72.7% of the variance while PC2 explains 3.2%. Regarding association to ancestry group, we found that PC1 P_ANOVA_ = 0.45 and PC2 P_ANOVA_ = 0.6 after correcting for batch year. Note that ATAC-seq read depth was the strongest contributing factor to the large variance in PC1.

### eQTL are concordant between ancestries

We first identified eQTLs within each ancestry group. Within The Netherlands cohort (N=150) we identified 3,133 eGenes at an FDR of 5% (Table S2), versus 1,394 eGenes in the Costa Rica cohort (N=46) (Table S3) and 1,492 eGenes in the Colombia cohort (N=50) (Table S4). Subsetting to matching SNP-gene pairs, we found an R2 of 0.93 between the Netherlands and Costa Rica eQTLs effects, an R2 of 0.96 between The Netherlands and Colombia eQTLs effects, and an R2 of 0.95 between the Colombia and Costa Rica eQTLs effects (Figure 2A). Combining genotype and expression data from the ancestry groups into one cohort (N=246) allowed us to perform a meta-analysis with greater power to detect associations, and minimal population stratification (lambdaGC = 1.08). This resulted in the identification of 5,258 eGenes at 5% FDR (Table S5).

**Figure 2.**
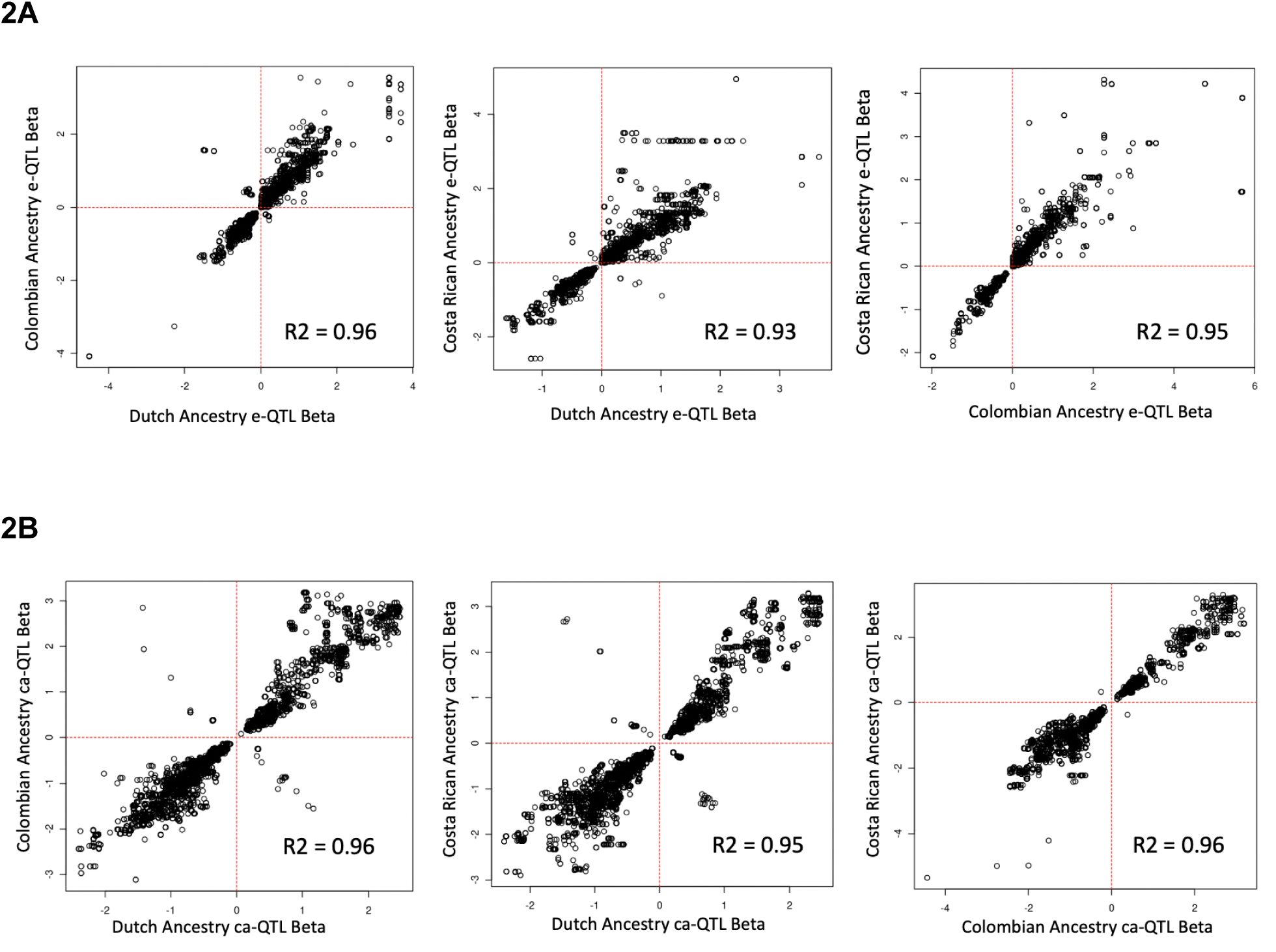
Expression and chromatin-accessibility QTLs are largely concordant between European and Latin American populations. **(A)** Each point represents a significant SNP-gene pair (note that many genes / peaks are associated with multiple SNPs, thus may be represented in multiple points). **(B)** Each point represents a significant SNP-peak pair.

### caQTL are concordant between ancestries

For chromatin-accessibility QTLs (caQTLs), we identified 2,001 peaks with FDR-significant association to genotypes in The Netherlands cohort (Table S6), 1,292 peaks in the Costa Rica cohort (Table S7), and 1,625 in the Colombia cohort (Table S8). Subsetting to matching SNP-peak pairs, we find an R2 of 0.95 between the Netherlands and Costa Rica caQTLs effects, an R2 of 0.96 between the Netherlands and Colombia caQTLs effects, and an R2 of 0.96 between the Colombia and Costa Rica caQTLs effects (Figure 2B).

Similar to the meta-analysis eQTL mapping, combining the genotype and chromatin data from the ancestry groups into one cohort (N=246) gave better power to detect associations (lambdaGC = 1.00), resulting in 3,557 ePeaks at an FDR threshold of 5% (Table S9).

Very few opposite-effect eQTLs or caQTLs were found amongst these pairwise comparisons, though these few instances are likely due to differences in LD blocks and allele frequencies between these populations. Consistency in overall direction of the effects and significance of these findings remain after pruning for SNPs in LD, both for eQTLs (Figure S4) and caQTLs (Figure S5).

### Causal effect correlations using local ancestry

To determine the degree of concordance between causal variant effect sizes across haplotype blocks, we used admix-kit^17^ to compute the genetic correlation across local ancestries, r_admix_, for both caQTLs and eQTLs. Prior to estimation of r_admix_, we utilized ADMIXTURE^18^ to determine ancestries of interest for our admixed individuals, which showed that the average admixture proportions of the Costa Rican and Colombian individuals were approximately 62% European, 33% Latin American, 5% West African, and 1% East Asian (Figure 3). Given that European and Latin American haplotypes made up the vast majority (>95%) of the ancestry admixture for these individuals, a two-way admixture was performed on 97 admixed individuals selected based on joint PCA with populations from the 1000 Genomes reference panel.

**Figure 3.**
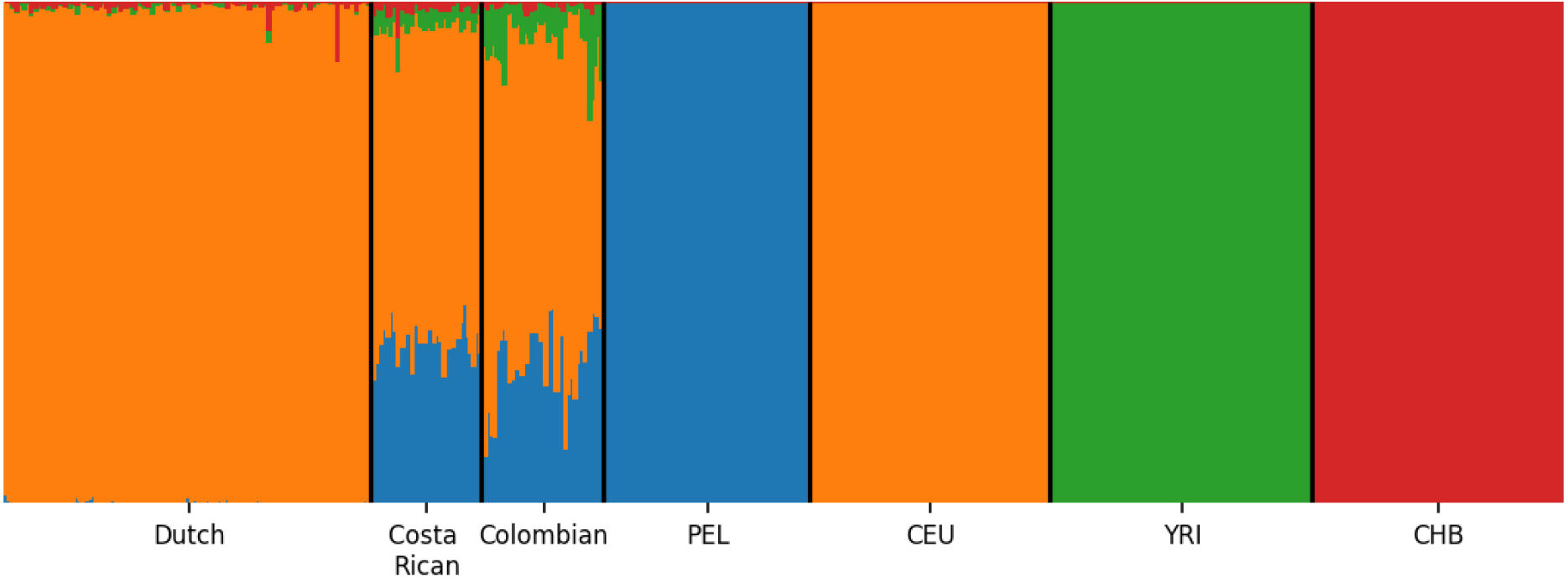
Proportions of reference populations via ADMIXTURE. Population reference panels from 1000 Genomes include PEL (Peruvian from Lima, Peru; AMR superpopulation), CEU (Utah residents from Northern and Western Europe; EUR superpopulation), YRI (Yoruba in Ibadan, Nigeria; AFR superpopulation), and CHB (Han Chinese in Beijing, China; EAS superpopulation).

Then, given our assumption that cis-SNPs are more likely to affect expression than trans-SNPs, we obtained r_admix_ estimates for 39,850 caQTLs and 11,523 eQTLs. After exclusion of genes or peaks with low standardized heritability (<2.0) and low confidence interval (CI) widths (<0.5), results for 1,000 eQTLs (Table S10) and 987 caQTLs (Table S11) were meta-analyzed to obtain r_admix_ = 0.983 (C.I: [0.97, 0.996], p-value = 0.019) for caQTLs and r_admix_ = 0.956 (C.I: [0.943, 0.968], p-value = 1e-12) for eQTLs. This suggests that the causal allelic effect sizes are the same across ancestries for these QTL.

### Fine-mapping of causal pathways

In order to identify potential causal paths starting from SNPs impacting the accessibility of a chromatin region which in turn leads to changes in expression of a proximal gene, we used the *pathfinder*^19^ framework. Briefly, *pathfinder* uses a hierarchical statistical framework to fine-map SNPs with chromatin marks and chromatin marks with gene expression in order to predict causal paths from SNP to mark to gene expression.

Given the paired-sample design of this analysis, we decided to continue only with the pooled analysis, rather than limit power by subsetting each ancestry group. We defined 100kb regions centered around the transcriptional start sites (TSS) of the 5,258 eGenes identified in the meta-analysis eQTL analysis, though we filtered these down to regions with prior evidence of gene-chromatin and chromatin-SNP associations (see **Methods**), resulting in a total of 934 gene regions. This low retention of regions is consistent with the empirical data analysis performed in the original *pathfinder* study (17.7% here and 8.9% in Roytman et. al^19^). Of these 934 tested gene regions, we found 203 genes with a posterior probability (PP) over 90% contained within the top ten paths in the region, and of these, 28 genes with PP>=90% in a single SNP-peak pair, suggesting high confidence that the expression of these genes is causally impacted by the fine-mapped SNP and chromatin mark (Table 1 for top paths, Table S12 for all paths).

We also investigated the spatial relationships between the SNPs, peaks, and genes in the top paths compared to a down-sampled set of random paths. Specifically, all paths with PP>=90% contained within the top 10 SNP/peak pairs for a gene were included as top paths, resulting in 1,893 paths. After randomly selecting 1,893 non-top paths, we plotted and compared the distributions of sizes between SNP-peak, SNP-gene, and peak-gene pairs (Figure S6). Two sample t-tests suggested no significant difference in SNP-peak distances between top paths and randomly generated paths (P=0.72), though significant differences were observed for SNP-gene (p=1.8e-15) and peak-gene (p<2.2e-16) distances. Top paths had significantly shorter distances between SNPs and genes compared to random paths. However, peak-gene distances were significantly longer in top paths compared to random paths, suggesting that these regions of open chromatin are enhancers to more distant genes.

### Transcriptome-wide associations to brain and skin-related traits

We performed a transcriptome-wide association study (TWAS) via the FUSION^20^ framework. Given the differences in ancestral background within the cohort, we split the analysis into European (Netherlands) and Latin American (Colombia and Costa Rica together) subsets. The ancestry-specific analysis resulted in 412 heritable eGenes from the European analysis, and 508 heritable eGenes from the Latin American analysis, with 67 genes overlapping between the two. Fisher’s exact test indicates this is a significant overlap (P = 6.2e-13), suggesting high confidence in the reproducibility across ancestries for these eGenes.

For association to GWAS risk loci, we tested summary statistics for bipolar disorder^7^ (multi-ancestry, though mostly European) and SCZ (both European^21^ and Latin American^22^ ancestry) as well as fibroblast-related traits including the UK BioBank^23^ dermatological disease traits eczema and psoriasis. We opted to include skin-related traits given that the functional data is collected from primary fibroblasts, and brain-related traits given the inclusion of individuals diagnosed with bipolar disorder. The association testing was performed separately per ancestry, then resulting Z scores were meta-analyzed via inverse variance weighting (IVW). Pooling data across ancestries has been shown to lead to false positives,^24^ though we chose to include this strategy to compare results across the different approaches.

For bipolar disorder, we identified two significant genes with P_IVW_ <= 1e-4 (Bonferroni-corrected for 500 heritable genes), including *LRP5L* (P_IVW_ = 2.4e-7) and *RPL27A* (P_IVW_ = 1e-5). A total of eight genes were significantly associated with the European-ancestry SCZ GWAS, including one gene shared with the BD-TWAS, *LRP5L* (P_IVW_ = 8.1e-8). Other genes identified using the European-ancestry SCZ GWAS included *GABARAP* (P_IVW_ = 9.9e-8), *LRRC37A4P* (P_IVW_ = 4.4e-7), *PLD2* (P_IVW_ = 1.5e-6), *RNF169* (P_IVW_ = 2.1e-6), *ILF2* (P_IVW_ = 2.5e-6), *TCHP* (P_IVW_ = 2.7e-5), and *PMM1* (P_IVW_ = 8.8e-5). *LRRC37A4P* was listed in the PGC schizophrenia study as an SMR-significant prioritized gene, based on PsychENCODE gene expression data. No genes were significantly associated with the Latin American-ancestry SCZ GWAS after Bonferoni correction (likely due to the small sample size of the GWAS). However, seven genes were nominally significant with the Latin American-ancestry GWAS, including *NCAM1* which was also nominally associated (0.05 > P > 1e-4) with the European ancestry SCZ and BD GWASes.

For the skin-related traits, we found three genes significantly associated with eczema, including *FBL* (P_IVW_ = 7.1e-5), *PMM1* (P_IVW_ = 2.1e-5) and *ILF2* (P_IVW_ = 9.0e-7), with the latter two interestingly also associated with schizophrenia in our TWAS analysis above. We did not find any Bonferroni-significant genes associated with psoriasis. See Figure 4A for TWAS results with all tested traits after IVW meta analysis.

**Figure 4.**
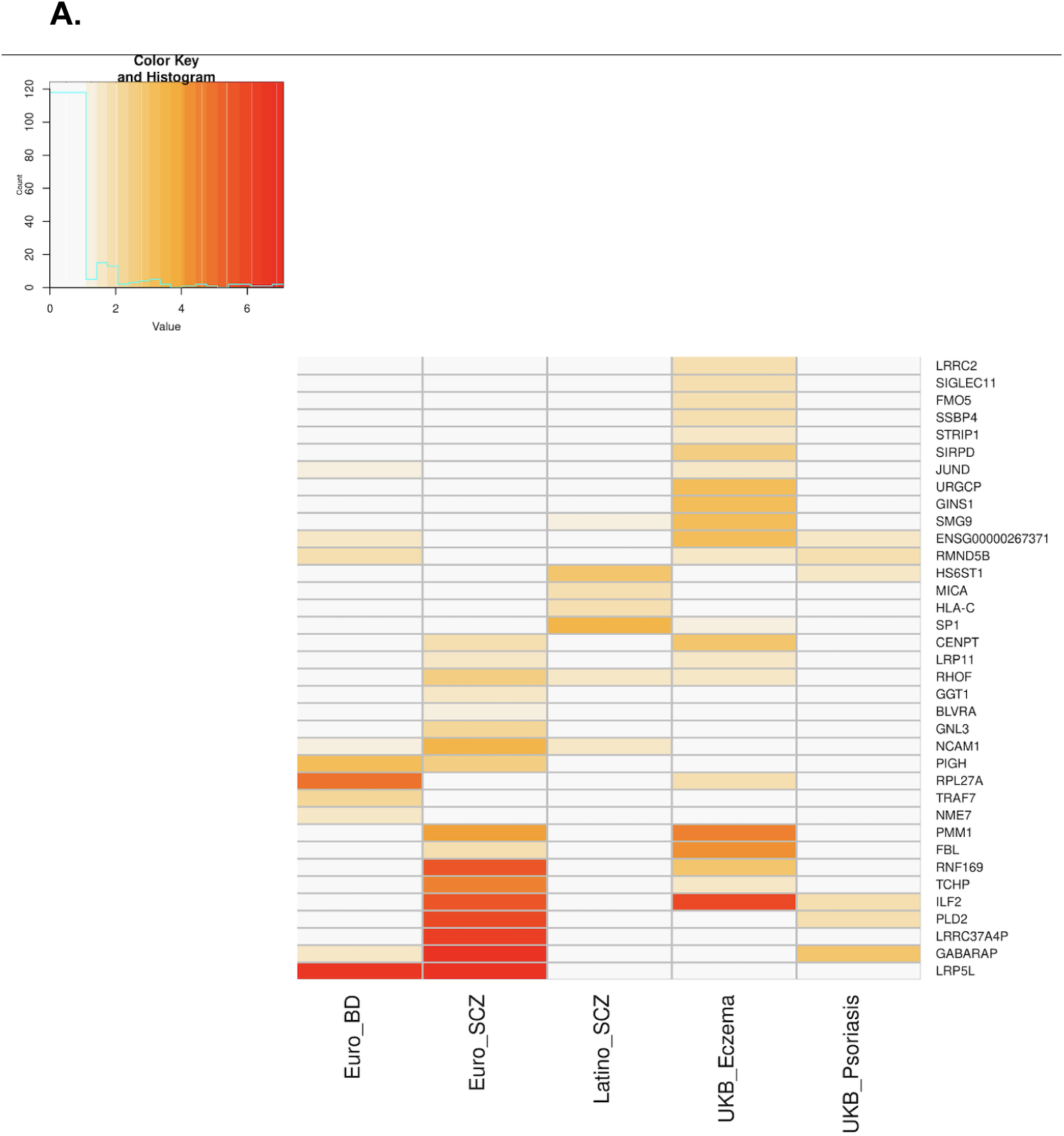
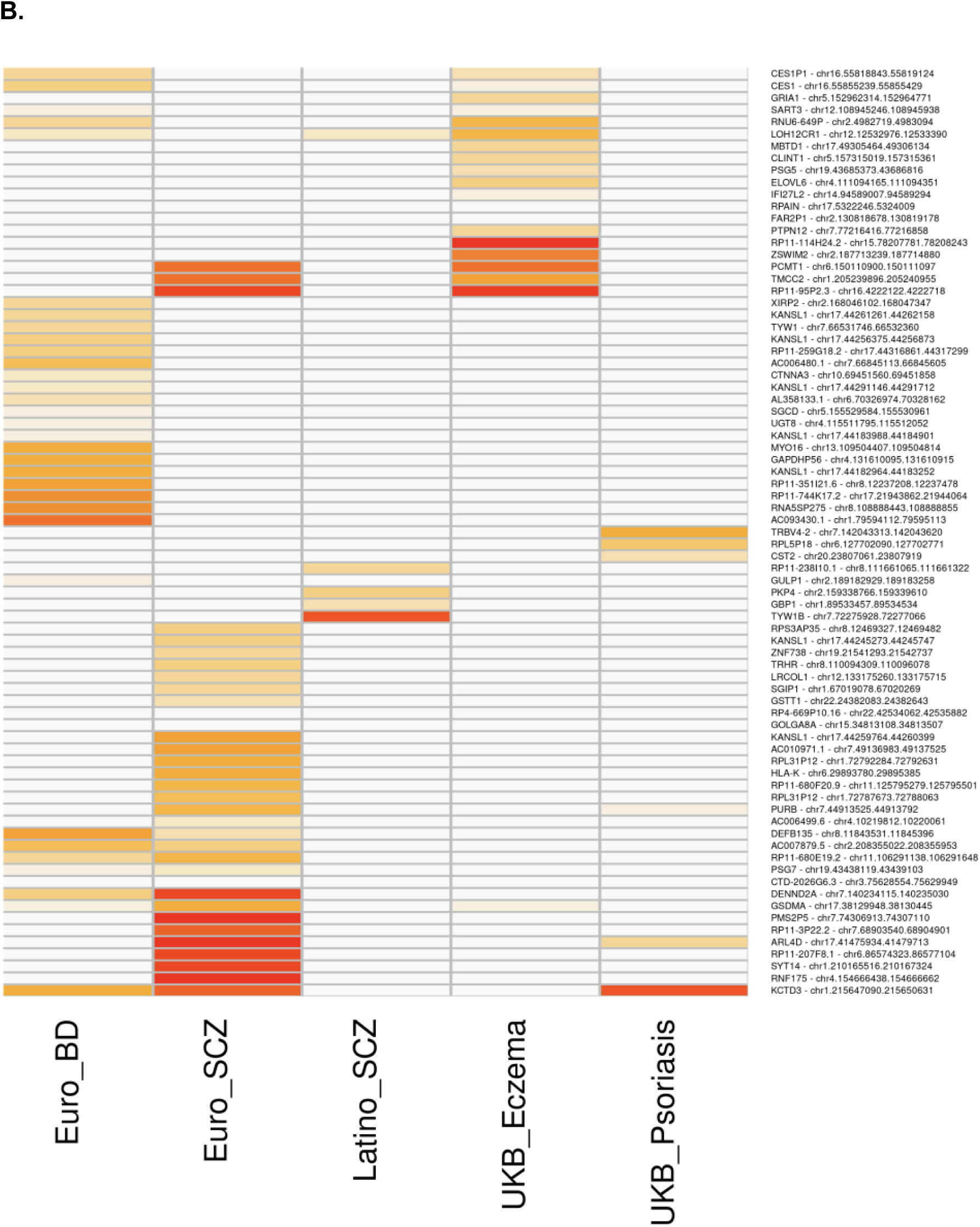
Transcriptome-wide (A) and chromatin-wide (B) association analyses with brain and skin-related traits. Values in the cells refer to −log10(p-value), with darker shades of red corresponding to lower p-values. Gene names are annotated for the rows of the TWAS heatmap. Rows of the CWAS heatmap are annotated with the most proximal gene and the open chromatin region boundaries.

When comparing the ancestry-specific results or the IVW-meta analysis results with the pooled-analysis TWAS, we found almost no significant eGenes in common for any traits, providing further evidence that pooled-ancestry analysis may be unreliable in the context of TWAS. Only *NCAM1* was significantly associated with European-ancestry SCZ both in the pooled analysis and IVW-meta analysis (though only nominally associated). See Table S13 and S14 for meta analysis and per-ancestry TWAS results, respectively, including SNP-based heritability for each gene.

### Chromatin-wide associations to brain and skin-related traits

We performed a chromatin-wide association study (CWAS,^10^ also called cis-trome wide) by again leveraging the FUSION-TWAS framework to associate regulatory elements with GWAS summary statistics. For association to GWAS risk loci, we used summary statistics for the same brain-related traits and dermatological traits as the TWAS analysis. Also paralleling the TWAS analysis, the association testing was performed separately per ancestry then meta-analyzed via inverse variance weighting (IVW).

We identified four P_IVW-_significant open chromatin regions with association to bipolar disorder (corresponding to four different proximal genes, see **Methods**). Two of these regions were proximal to brain-expressed genes (*SYT14* and *SGIP1*). We identified 16 significantly associated open chromatin regions for European-ancestry schizophrenia (across 11 proximal genes); and no associated open chromatin regions for Latin American-ancestry schizophrenia. Of the genes that we identified to be proximal to these open chromatin regions, only *KANSL1* (corresponding to open chromatin region chr17:44182964-44183252bp) has shown prior evidence of association to schizophrenia, specifically using fetal brain gene expression from PsychENCODE. For the skin-related traits, we found 12 open chromatin regions (across 12 proximal genes) significantly associated with psoriasis, and 24 chromatin regions significantly associated with eczema (across 23 proximal genes). See Figure 4B for CWAS results after IVW meta analysis. See Table S15 for meta-analysis results and S16 for per-ancestry results, including peak heritability.

## DISCUSSION

In this study, we performed QTL analysis on gene expression and chromatin accessibility data from human primary fibroblasts of a multi-ancestry bipolar disorder sample set. QTL effect sizes were found to be concordant between the European and admixed American populations, suggesting that the genetic variants that affect either gene expression or chromatin accessibility tend to have similar effects across these populations. However, while the effect sizes may be consistent, the underlying genetic architecture and linkage disequilibrium (LD) patterns can vary significantly between ancestries,^25^ contributing to the clustering of ancestral groups in principal component analysis of genetic data. For more reliable integration of molecular data into GWAS, it is necessary to match the genetic ancestral background to functional omics datasets and LD reference panels. This approach not only improves the accuracy of genomic studies^25,1^ but also fosters inclusivity and furthers genomic research efforts on a global scale.

Multi-omics datasets have advanced the field of molecular biology by providing a comprehensive and integrated view of biological systems at various levels. By collecting genomics, transcriptomics, and chromatin data across hundreds of individuals, we and others have shown that such datasets can unravel molecular mechanisms underlying biological processes. Such comprehensive analyses facilitate in deciphering complex biological patterns and predicting interactions at the molecular level. Regarding our TWAS and CWAS analyses, we find more than twice the number of chromatin peaks (87) than genes (36) associated with the tested traits. This is consistent with the prostate cancer findings in the original CWAS method paper,^10^ providing further evidence to the idea that gene expression tends to be more context-dependent than chromatin accessibility^14^ and that GWAS loci tend to be depleted of eQTLs.^11^

As an illustrative example of the merits of combining multi-omics data from a single cohort, we were able to finemap a possible pathway for the *MICA* gene (major histocompatibility complex (MHC) class I polypeptide-related sequence A). *MICA* has been previously found to be associated with schizophrenia,^26,27^ and integrating the expression data for this gene region with chromatin data suggested a causal pathway with posterior probability >0.98 from rs2442724 to a locus of open chromatin in the MHC region (chr6:31,367,110-31,369,941), upstream of the *MICA* gene (chr6:31,368,488-31,383,092). While the MHC region is known to be difficult to parse given extensive LD,^28^ the large cohort of paired transcriptome/cistrome data allowed for fine-mapping of the regulation of this particular gene. Several MHC genes have been previously reported as associated with psychiatric disorders,^29,30–32^ implicating this region as a probable risk locus.

Another gene, Long Intergenic Non-Protein Coding RNA 933 (ID: ENSG00000259728, genomic location chr15:85,114,155-85,121,355), for which we identified a highly probable (PP=0.99) pathway has been previously associated with BD^33^ and SCZ^21^ via the PsychENCODE TWAS.^34^ The causal path included SNP rs12900391 with a chromatin peak at chr15:84,542,517-84,544,020. While the exact function of this gene is not well understood, the genetic associations with BD found by the latest wave of the Psychiatric Genomics Consortium have suggested the involvement of pathways regulating insulin secretion, calcium channel activity, and signaling of endocannabinoids and glutamate receptors.^33^ We also found a probable pathway for the gene Gamma-Aminobutyric Acid (GABA) Receptor Subunit Rho-2 (*GABRR2)*. GABA is the major inhibitory neurotransmitter in all mammalian brains, and while direct association between BD or SCZ and this gene has not yet been detected, the locus of this gene on chromosome 6q has had associations to psychiatric illness.^35–37^ Similarly, we identified a probable pathway for the *GATAD2A* gene, previously found to be associated with SCZ in the latest GWAS.^21^ We identified probable pathways for other interesting brain-relevant genes including *CRCP*, which encodes a membrane protein that functions as part of a receptor complex for a small neuropeptide that increases intracellular cAMP levels in the brain.^38^

While we identify novel molecular mechanisms for genes potentially relevant to disease biology, there are several limitations to our study. Although fibroblasts can offer valuable insights into certain biological processes due to shared cis-genetic effects, their relevance to complex brain-specific pathways remains limited. However, fibroblasts have previously been used in studying circadian rhythms, a phenotype which is known to be dysregulated in bipolar disorder, suggesting that relevant molecular mechanisms are at least partially preserved.^39,40^

Secondly, this investigation focused solely on cis-QTLs, while trans-QTLS, though more difficult to ascertain, could potentially unveil additional regulatory elements involved in the brain and skin-related traits studied here. Furthermore, while our study is an important step forward in the inclusion of Latin American individuals in genomic studies, the lack of samples from individuals of African and Asian countries and other diverse ancestries represents a limitation, as genetic variants that are common to these populations may be missing or very rare in the European and Latin American samples included here. Lastly, the scarcity of Latin American-based GWAS datasets on which to perform ancestry-matched TWAS underscores the urgent need for more extensive efforts to incorporate diverse ancestral backgrounds in genetic studies. Utilizing GWAS and functional data from diverse ancestries is crucial for understanding the genetic basis of complex traits in a globally inclusive manner.

## METHODS

### Sequencing data collection

Skin biopsies were collected in 2010-2012 from individuals of Colombian ancestry and Costa Rican ancestry, and collected in 2013-2014 from individuals of Dutch ancestry to generate human primary fibroblasts. Primary fibroblast cultures were established following standard procedures^41^ and stored as frozen aliquots in liquid nitrogen. Fibroblasts were thawed out in batches of 8 lines at a time and grown to confluence in T75 culture flasks in standard culture media (DMEM containing 10% fetal bovine serum (FBS) and 1x Penicillin-Streptomycin). Upon reaching confluence, cells were passaged to 12 well plates at a density of 1×10^5^ for ATAC and 2×10^5^ for RNA. The next day cells were collected for further processing.

This cohort includes BD patients and unrelated healthy controls. DNA and RNA extractions from these cells were performed simultaneously from the same batch of cells in order to minimize technical artifacts that may confound the mediation analysis. Relatedly, nine of the Netherlands samples were sequenced for RNA-seq and ATAC-seq in both batches and correlation of the gene expression and peak intensities was high, particularly for gene expression (Table S17).

For duplicated sample pairs with high correlation (R2>0.85) for both gene expression and chromatin peak intensity, we randomly selected one in the pair of IDs to include in downstream analyses. For the two pairs with low correlation in chromatin peak intensities, quality control revealed that the second batch had significantly lower read depth for these individuals, thus we included the samples originating from the first batch in these instances.

### RNA-seq data generation and processing

RNA was extracted from fibroblasts in order to assess the levels of gene expression across the genome. Cells were lysed using 350uL RLT lysis buffer from the Qiagen RNeasy mini kit. Lysed cells were then scraped off the plate, transferred to a Qiaschredder (Qiagen 79656) and centrifuged for 2 min at max speed to further homogenize. Cell lysates were kept in −80 until extraction. RNA from cell lysates was extracted using the Qiagen RNeasy mini kit (Qiagen 74106). Cell lysates were extracted in a randomized order to prevent batch effects in downstream analysis. In order to collect total RNA including small RNAs, the standard extraction protocol (Purification of Total RNA from Animal Cells using Spin Technology) was adjusted by making the following changes:

- adding 1.5 volumes of 100% ethanol, instead of 70%, after the lysis step (step 4 in handbook protocol)
- adding 700 mL of buffer RWT (Qiagen 1067933) instead of the provided RW1 (step 6 in handbook protocol)

TruSeq Stranded polyA selected library preps were generated and samples were sequenced on the Illumina HiSeq 4000 sequencer with 75-base paired end reads, at an average of 50 million mapped reads per sample. The resulting FASTQ files were pseudo-aligned to hg19 using kallisto,^42^ resulting in a matrix of transcripts per million (TPMs) which were aggregated to the gene level. The expression matrix was filtered for protein-coding genes and outliers based on technical variation (n genes = 18,886 remain after filters). Principal component analysis was initially performed on the log-transformed counts matrix and resulting PCs were assessed for correlation with batch year and ancestry. For downstream analyses, we corrected the expression matrix by including covariates for sex, first two genotype PCs, and the year of the sequencing batch (for Netherlands individuals). Previous studies have shown that there are likely unmeasured or “hidden” factors that reduce the power to detect associations in next-generation sequencing data, therefore we also performed PEER^43^ (Probabilistic Estimation of Expression Residuals) factor analysis to find such hidden determinants of variation. PEER factors were computed separately per ancestry group, including 5 factors for the Colombian and Costa Rican groups, and 15 for the Netherlands group, given the difference in sample sizes. The resulting residuals matrix that remained after accounting for all factors was then used as input for downstream analyses.

### ATAC-seq data generation and processing

ATAC-seq libraries were generated as previously described.^44^ Samples were sequenced on the Illumina HiSeq 4000 sequencer with 75-base paired end reads, at an average of 39 million mapped reads per sample. Trimmed reads were aligned to the hg19 reference genome using bowtie2^45^, and filtering steps were taken to remove unmapped reads, non-primary alignment, and low-quality reads, as recommended by the ENCODE standards for ATAC-seq data analysis.^46^ Reads were then input into MACS2^47^ in order to call peaks that are significantly enriched against the local background using a false-discovery rate (FDR) correction threshold of 0.05 and modeled by the Poisson distribution. From this initial set of peaks, blacklisted regions^48^ as identified by ENCODE were removed in order to exclude regions that have anomalous or unusually high signals due to repetitive or unstructured sequences. Then, the remaining peaks across all samples were combined to form the consensus regions, using the bedtools intersect function to stitch together any regions within 147bp into one larger peak region. This resulted in over 418,000 consensus peaks, which were limited to only those peaks that are called in at least 30% of individuals, reducing the number of peaks to 77,957. These consensus regions, in conjunction with the filtered reads (bam files) per individual, were used in the R package featureCounts to determine the number of reads each individual had within each peak, with higher counts of reads per peak indicating greater accessibility of that region of the genome.

This resulted in an *N* x *P* peak counts matrix where *N* = number of individuals and *P* = number of peak regions. This matrix was log-transformed to account for skewness and ensure normalization. Principal component analysis (PCA) was then performed on the log-transformed counts matrix to assess for batch-specific and ancestry-specific correlations with PCs. PCs were correlated via Spearman’s rank correlation against other technical factors in order to determine drivers of variation within the data that are unrelated to underlying biology. Using a Bonferroni significance level of P < 2e-05, we found that sex, read depth, fraction of mitochondrial DNA, TSS enrichment score, median fragment size, and fraction of reads in peaks were correlated with the first two ATAC-seq PCs. The sequencing batch year was also included in order to account for batch effects.

Using the log-transformed counts matrix as the input measures and the independent technical factors identified from the PC correlations plus age as covariates, we used PEER to find hidden confounders. PEER factors were computed separately per ancestry group, including 5 factors for the Colombian and Costa Rican groups, and 15 for the Netherlands group, given the difference in sample sizes. The resulting residuals matrix that remained after accounting for all factors was then used as input for the QTL analysis.

### Genotyping and imputation

Samples were genotyped and imputed separately in two batches, the first consisting of 129 individuals of European ancestry genotyped via the OmniExpressExome platform, with the second batch consisting of 21 individuals of Dutch ancestry, 50 individuals of Colombian ancestry, and 46 individuals of Costa Rican ancestry genotyped via the Global Screening Array. Genotypes were first filtered for Hardy-Weinberg equilibrium p value < 1.0e-6 for controls and p value < 1.0e-10 for cases, with minor allele frequency (MAF) > 0.01. Genotypes were then imputed into the 1000 Genomes Project phase 3^49^ reference panel by chromosome using RICOPILI v.1^50^ separately per genotyping platform, then subsequently merged, applying an individual-missingness threshold of 10%, SNP-missingness of 5%, and MAF > 0.05 for post-merge quality control. Imputation quality was assessed by filtering variants where genotype probability > 0.8 and INFO score > 0.1 resulting in 2,747,786 autosomal SNPs that were common across both datasets.

### QTL mapping

QTL analysis was performed with MatrixEQTL^51^ using a cis-locus distance defined as +/-1Mb around the peak midpoint or gene TSS, initially done separately per ancestry group. We included the identity by state (IBS) similarity matrix of the genotypes as an error covariance term within the model. Associations that remain after an FDR threshold of 5% were retained for downstream analysis. Correlations above R2 of 90% of the resulting SNP-gene or SNP-peak effect sizes per ancestry group suggested that the groups could be combined for a gain in power, and thus the analysis was repeated with all individuals together and with an added covariate term for ancestry.

### Assessing ancestry-specific causal effects

Ancestry specific causal effects were calculated using admix-kit^17^ on 97 admixed individuals, including 45 Costa Ricans, 50 Colombians, and 2 Dutch. These admixed individuals were determined by performing a joint PCA with the imputed genotype and 1000 Genomes Project reference panel.^49^ The first 4 PCs, maximum sample distance of 1.5, sample t-range of (0.05, 0.95), and super populations EUR and AMR were used via the select-admix-indiv function in admix to select individuals to include in our analysis. To determine the super populations as well as the ancestries to use for the correlation estimates, we utilized ADMIXTURE^18^ to compare our cohort with four reference populations, including Europeans (CEU), West Africans (YRI), Latin Americans (PEL), and East Asians (CHB) from 1000 Genomes Project to determine admixture proportions with supervised analysis. With these admixed individuals, we inferred local ancestry using RFmix.^52^

The admix-kit package was used to determine the similarity among the genome-wide causal allelic effects across specified local ancestries in admixed individuals.^17^ To calculate genetic correlations for each gene or peak, we built a window-based genetic relationship matrix, GRM, using a cis-locus distance defined as +/- 1Mb around each peak or gene rather than a genome-wide GRM to focus on the contribution of cis-SNPs on expression. The window-based GRM is used to estimate log-likelihood at different r_admix_ values to obtain the point estimate, credible interval, and p-value for each gene or peak. To obtain the genetic correlation across all genes or peaks, we meta-analyze across the genes/peaks via the meta-analyze-genet-cor() function in admix.

Prior to meta-analysis, due to our low sample size, we exclude genes or peaks whose standardized heritability (hsq_est/hsq_stderr) is less than 2, confidence interval widths are below 0.5, and genes/peaks that had more than one credible interval.

### Fine-mapping of SNP-chromatin-gene pathways

We used the *pathfinder* method which accounts for both SNP LD and the correlation structure between chromatin marks by using a multivariate normal distribution. This method iterates through each possible path to determine its corresponding posterior probability, thus enabling us to prioritize SNPs and chromatin peaks that mediate gene expression, which can then be prioritized for functional validation. We restricted the regions tested by taking the transcriptional start site (TSS) for each eGene and pulling out all SNPs and chromatin peaks within 50kb upstream or downstream of the TSS. These regions were then filtered via a two-stage regression analysis, wherein the gene expression values were regressed on the proximal chromatin marks, and for models with a resulting p-value less than 0.05, we regressed SNP genotypes in that region onto the residuals from the initial peak-gene regression. Any regions with at least one p-value less than 0.05/(number of SNPs in region) were retained for pathfinder analysis. All chromatin peaks within the region were correlated pairwise via the cor() function in R, and LD for all SNPs within the region was calculated through plink –r2 square. We then inputted these regions into the pathfinder.R script (https://github.com/meganroytman/pathfinder/blob/master/pathfinder.R).

### TWAS

To identify eQTL associated with GWAS traits, we performed a TWAS using the FUSION^20^ software (http://gusevlab.org/projects/fusion). First, we generated weights for all 5,258 FDR-significant eGenes using the FUSION.compute_weights.R script, restricted to loci +/- 1Mb around the lead SNPs per each gene. We used the PEER-corrected gene expression thus no additional covariates were included in the model. In generating the weights, eGenes were first filtered for those with significant SNP-heritability (p-value <= 0.05).

We used the FUSION.assoc_test.R script to test for association between the gene weights and GWAS for bipolar disorder, schizophrenia (EUR and AMR ancestry), and UKBB skin-related GWAS for eczema and psoriasis. Gene-trait pairs were selected based on the best performing model after five-fold cross validation, including for Best Unbiased Linear Predictor (BLUP), elastic net (ENET), Least Absolute Shrinkage and Selection Operator (LASSO), and just using the top SNP. To account for LD structure we used an in-sample LD panel. The --coloc flag was included to perform colocalization^53^ on any genes that had an association with the trait of interest with TWAS.P < 0.05.

### CWAS

Similarly, to identify caQTL associated with GWAS traits, we performed a CWAS^10^ using the FUSION software. First, we generated weights for all FDR-significant ePeaks using the FUSION.compute_weights.R script, restricted to loci +/- 1Mb around the lead SNPs per each gene. We used the PEER-corrected peak intensities thus no additional covariates were included in the model.

We used the FUSION.assoc_test.R script to test for association between the peak weights and GWAS for bipolar disorder, schizophrenia (EUR and AMR ancestry), and UKBB skin-related GWAS for eczema and psoriasis. Peak-trait pairs were selected based on the best performing model after five-fold cross validation, including for Best Unbiased Linear Predictor (BLUP), elastic net (ENET), Least Absolute Shrinkage and Selection Operator (LASSO), and just using the top SNP. To account for LD structure we used an in-sample LD panel. The --coloc flag was included to perform colocalization on any peaks that had an association with the trait of interest with TWAS.P < 0.05. Proximal genes were identified using the GenomicRanges^54^ package in R, employing the nearest() function.

## Supplemental Figures and Table Captions

**Figure S1: Principal component analysis of sample cohort overlaid onto 1KG.** EUR = European, AFR = African, EAS = East Asian, SAS = South Asian, AMR = American, SAMPLE = cohort analyzed in this study.

**Figure S2: Principal component analysis of gene expression data after correction for covariates.** Annotated by country of origin on the left and year of sequencing batch on the right.

**Figure S3: Principal component analysis of chromatin accessibility data after correction for covariates.** Annotated by country of origin on the left and year of sequencing batch on the right.

**Figure S4: Pairwise comparison of effect sizes between ancestry-specific eQTLs.** Only including LD-independent SNPs

**Figure S5: Pairwise comparison of effect sizes between ancestry-specific caQTLs.** Only including LD-independent.

**Figure S6: Spatial relationships between SNPs, peaks, and genes in top vs random paths.**A. SNP-peak distances; B. SNP-gene distances; and C. Peak-Gene distances.

**Table 1: Genes with posterior probability of paths >=90% in a single SNP-peak pair.** Peaks are formatted as chr:startBP-endBP.

**Table S1: Variance explained and Analysis of Variance (AOV) p values for first 5 principal components of genotype, expression, and chromatin data with ancestry and sequencing year.**

**Table S2: Top eQTLs for Netherlands cohort, including SNP, ENSG, p-value and beta effect size.**

**Table S3: Top eQTLs for Costa Rica cohort, including SNP, ENSG, p-value and beta effect size.**

**Table S4: Top eQTLs for Colombia cohort, including SNP, ENSG, p-value and beta effect size.**

**Table S5: Pooled-cohort eQTL results.**

**Table S6: Top caQTLs for Netherlands cohort, including SNP, peak boundaries, p-value and beta effect size.**

**Table S7: Top caQTLs for Costa Rica cohort, including SNP, peak boundaries, p-value and beta effect size.**

**Table S8: Top caQTLs for Colombia cohort, including SNP, peak boundaries, p-value and beta effect size.**

**Table S9: Pooled-cohort caQTL results.**

**Table S10: Admixture results for tested genes. rg_mode refers to the genetic correlation between the local ancestry haplotypes of the eQTL; conf_inf refers to the confidence interval around the rg_mode values.**

**Table S11: Admixture results for tested peaks. rg_mode refers to the genetic correlation between the local ancestry haplotypes of the caQTL; conf_inf refers to the confidence interval around the rg_mode values.**

**Table S12: All paths for genes having at least 90% PP within its top 10 SNP/peak pairs.** Peaks are formatted as chr:startBP-endBP. Posteriors refer to posterior probability.

**Table S13: Meta-analysis of TWAS results for brain and skin-related traits.** Beta = IVW-meta analysis P value; se = IVW-meta analysis standard error; z = IVW-meta analysis z score; p = IVW-meta analysis p value; SumStats = cohort/trait summary statistics used.

**Table S14: TWAS-FUSION results for ancestry-specific and pooled analysis.** Abbreviations: CHR; chromosome, P0; gene start, P1; gene end, HSQ; heritability of the gene, BEST.GWAS.ID; rsID of the most significant SNP in locus, BEST.GWAS.Z; Z-score of the most significant GWAS SNP in locus, EQTL.ID; rsID of the best eQTL in the locus, EQTL.R2; cross-validation R2 of the best eQTL in the locus, EQTL.Z; Z-score of the best eQTL in the locus, EQTL.GWAS.Z; GWAS Z-score for this eQTL, NSNP; number of SNPs in the locus, MODELCV.R2; cross-validation R2 of the best performing model, MODELCV.PV; cross-validation P-value of the best performing model, SNP; single nucleotide polymorphism, FDR; false-discovery rate, eQTL; expression quantitative trait locus. PP; posterior probability (0: no association, 1: eQTL association only, 2: trait association only, 3: independent eQTL/trait association, 4: colocalized association); SumStats = cohort/trait summary statistics used.

**Table S15: Meta-analysis of CWAS results for brain and skin-related traits.** Beta = IVW-meta analysis P value; se = IVW-meta analysis standard error; z = IVW-meta analysis z score; p = IVW-meta analysis p value; SumStats = cohort/trait summary statistics used; Closest.Gene = gene symbol for nearest gene to chromatin region.

**Table S16: CWAS-FUSION results for ancestry-specific and pooled analysis.** Abbreviations: CHR; chromosome, P0; chromatin start, P1; chromatin end, HSQ; heritability of the peak, BEST.GWAS.ID; rsID of the most significant SNP in locus, BEST.GWAS.Z; Z-score of the most significant GWAS SNP in locus, EQTL.ID; rsID of the best caQTL in the locus, EQTL.R2; cross-validation R2 of the best caQTL in the locus, EQTL.Z; Z-score of the best caQTL in the locus, EQTL.GWAS.Z; GWAS Z-score for this caQTL, NSNP; number of SNPs in the locus, MODELCV.R2; cross-validation R2 of the best performing model, MODELCV.PV; cross-validation P-value of the best performing model, SNP; single nucleotide polymorphism, FDR; false-discovery rate, caQTL; chromatin accessibility quantitative trait locus. PP; posterior probability (0: no association, 1: caQTL association only, 2: trait association only, 3: independent caQTL/trait association, 4: colocalized association); SumStats = cohort/trait summary statistics used; Closest.Gene = gene symbol for nearest gene to chromatin region.

**Table S17: Duplicated Netherlands sample correlation for gene expression and chromatin accessibility data.** RNA R2 = correlation between gene expression TPMs of the two samples; ATAC R2 = correlation between chromatin accessible peaks of the two samples.

