## Supplemental Figures and Tables for "Quantitative trait loci mapping of gene expression and chromatin accessibility in primary fibroblast reveals shared allelic effects between Latin American and European ancestries": Figure3_admixture_proportions.docx

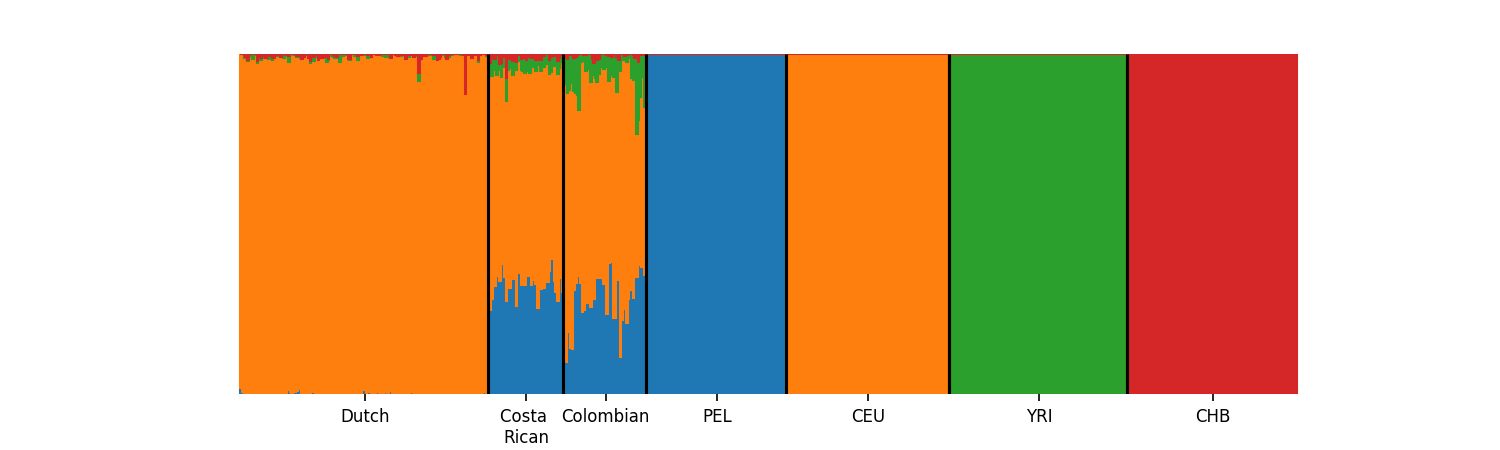

**Figure 3.** **Proportions of reference populations via ADMIXTURE.**

Population reference panels from 1000 Genomes include PEL (Peruvian from Lima, Peru; AMR superpopulation), CEU (Utah residents from Northern and Western Europe; EUR superpopulation), YRI (Yoruba in Ibadan, Nigera; AFR superpopulation), and CHB (Han Chinese in Bejing, China; EAS superpopulation).
