## Supplementary figures and images for "Quantitative trait loci mapping of gene expression and chromatin accessibility in primary fibroblast reveals shared allelic effects between Latin American and European ancestries"

### Figure4A_TWAS.tiff

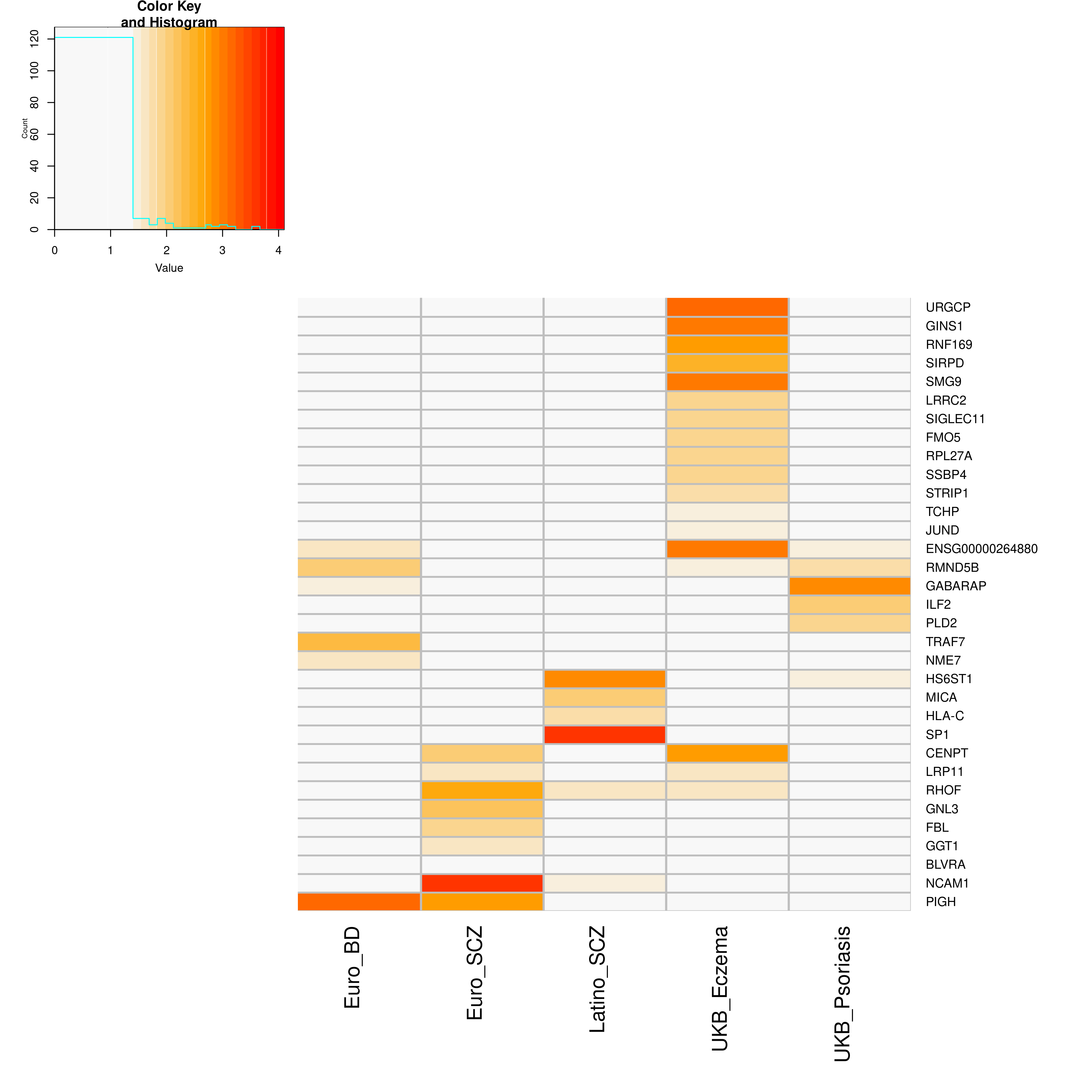

### Figure4B_CWAS.tiff

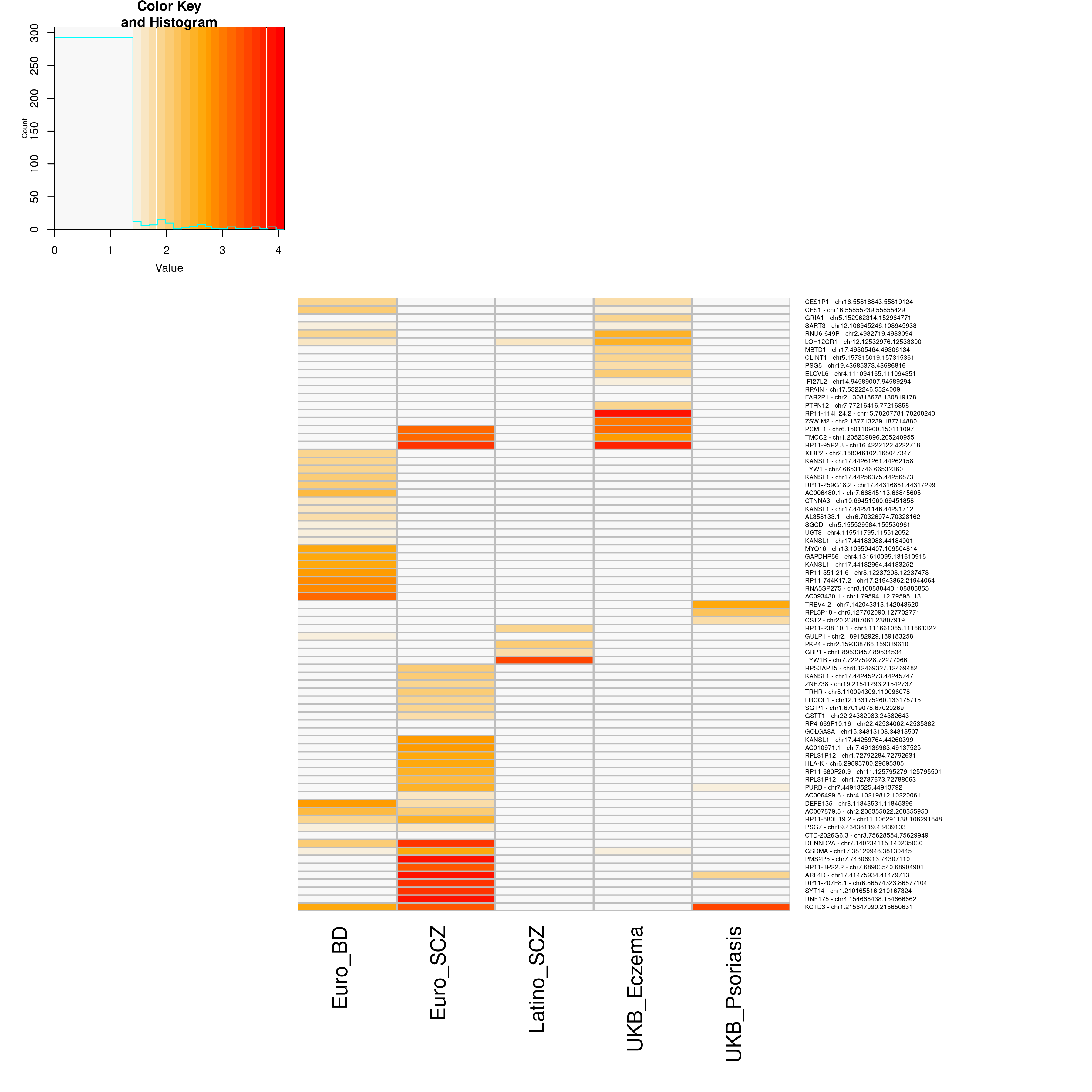

### FigureS1_PCA_with_1KG.docx

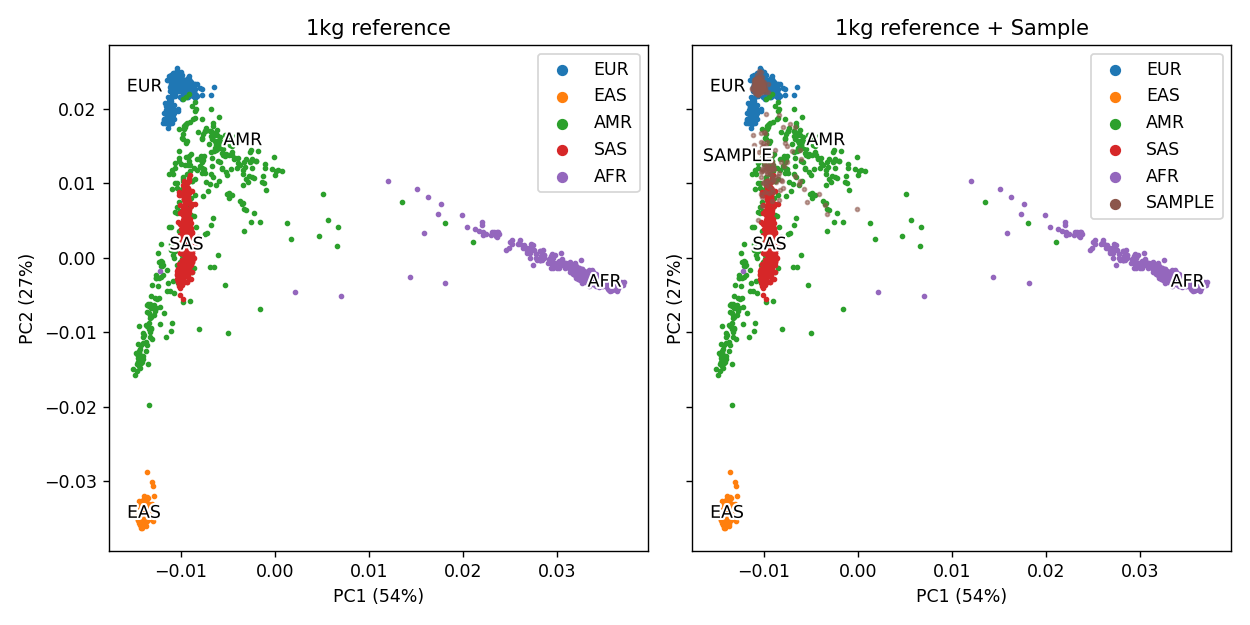

### FigureS2.png

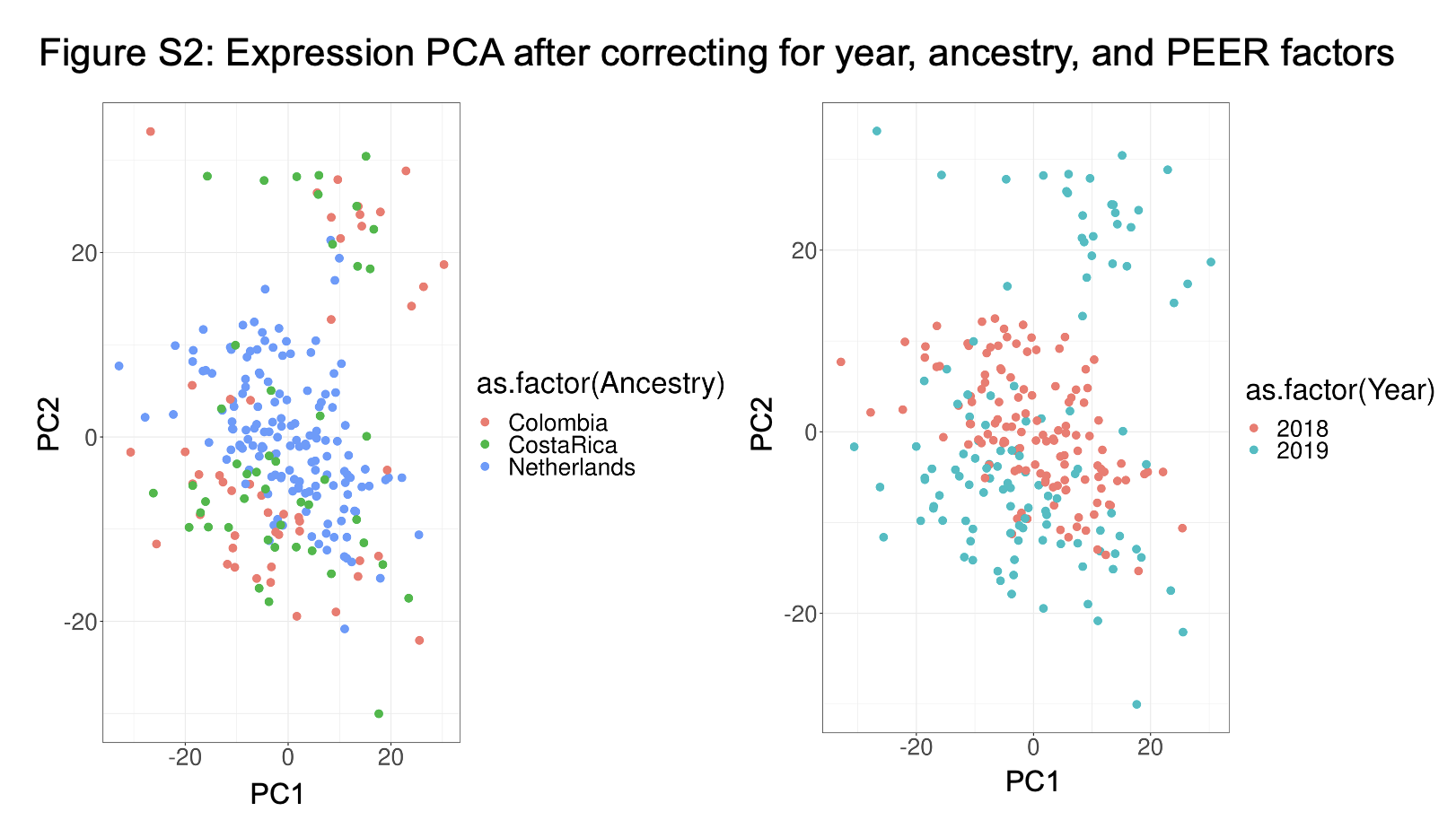

### FigureS3.png

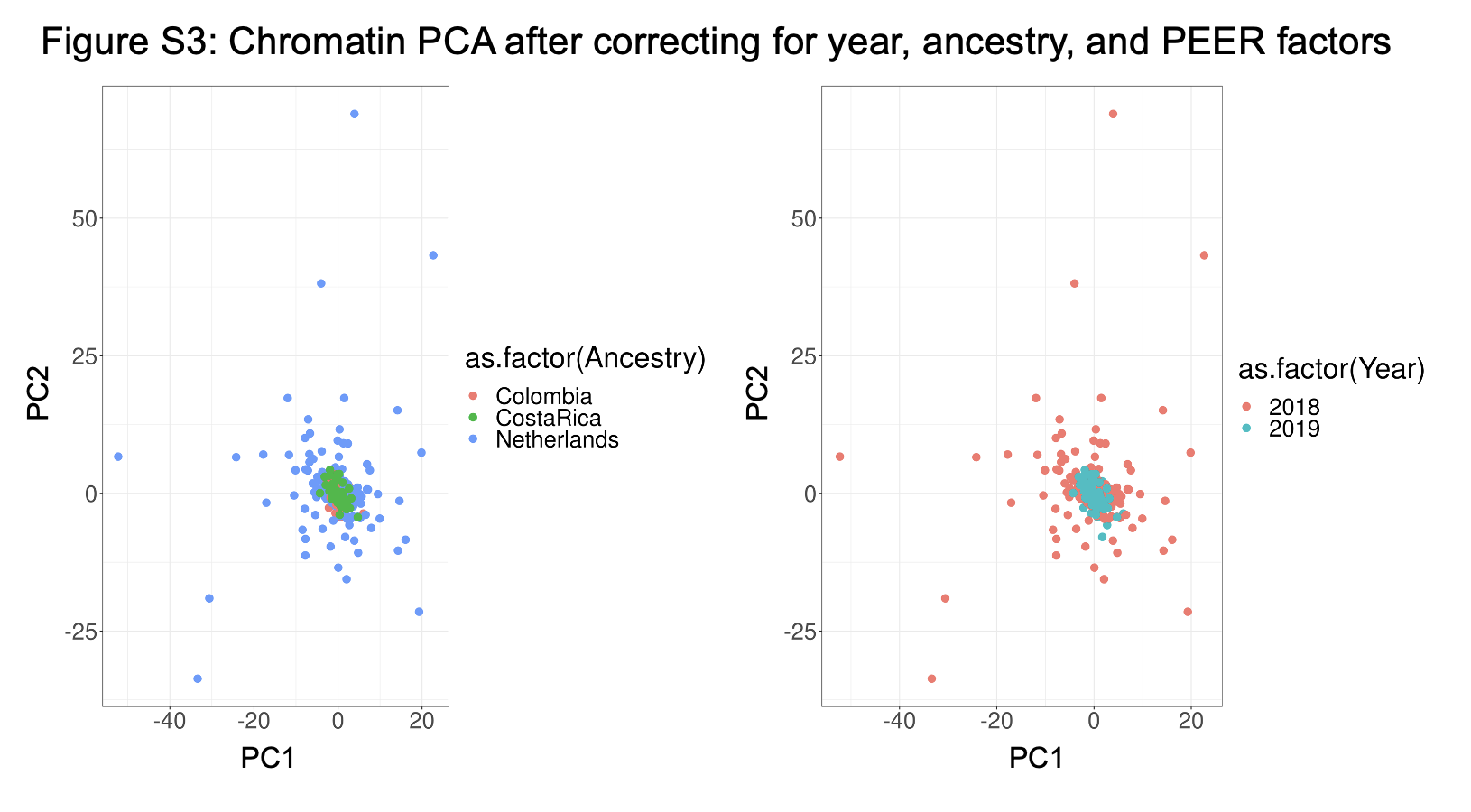

### FigureS4.png

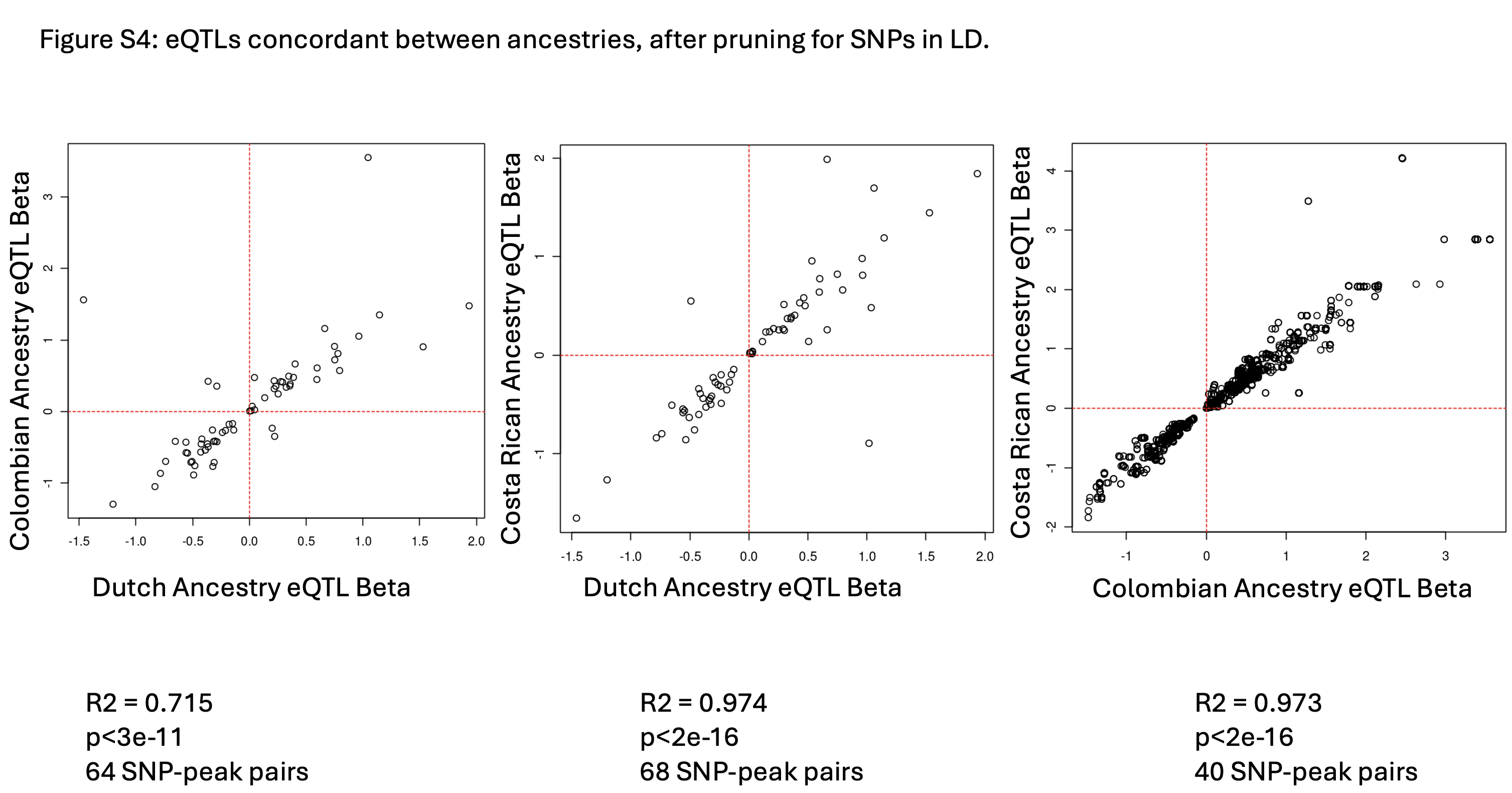

### FigureS5.png

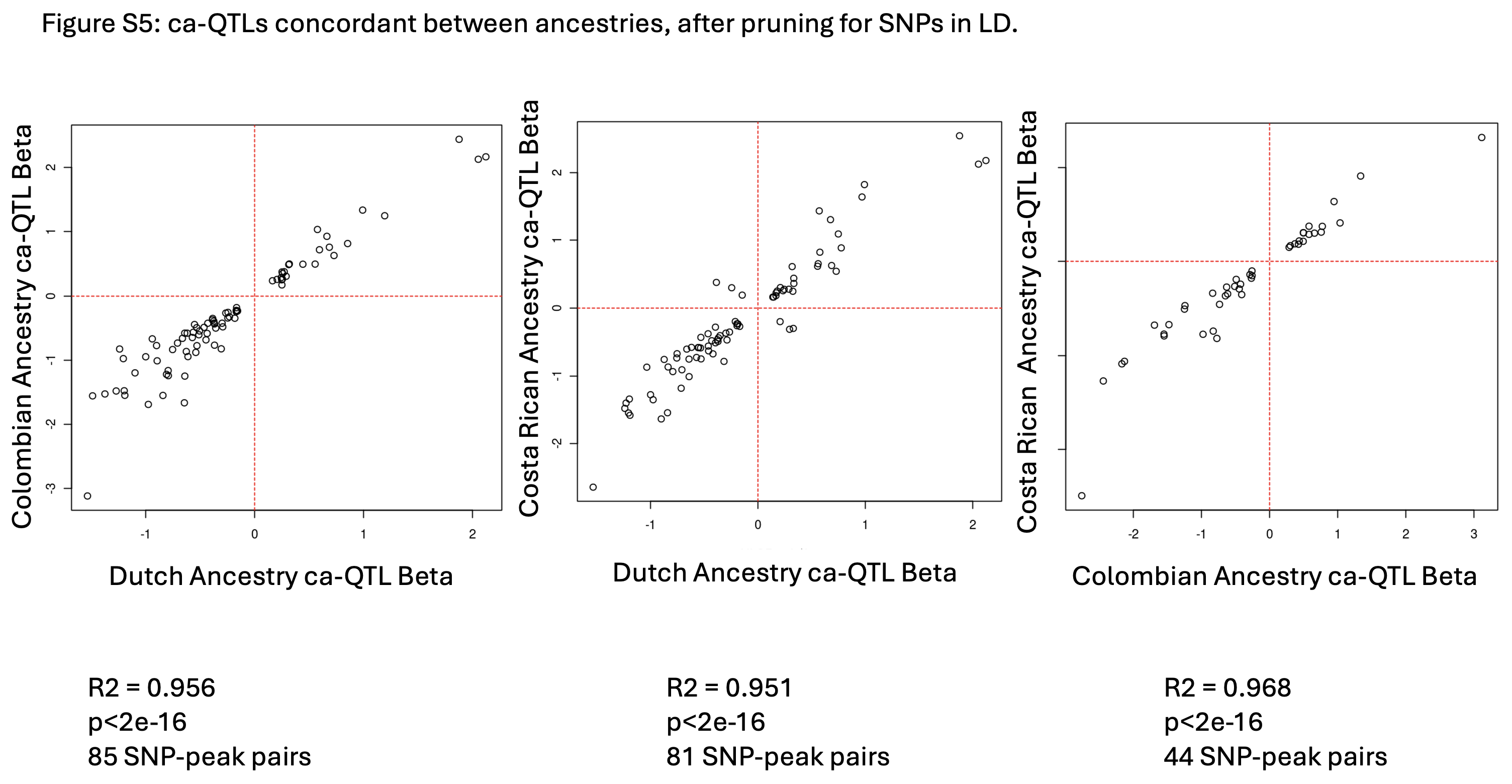

### FigureS6.pdf

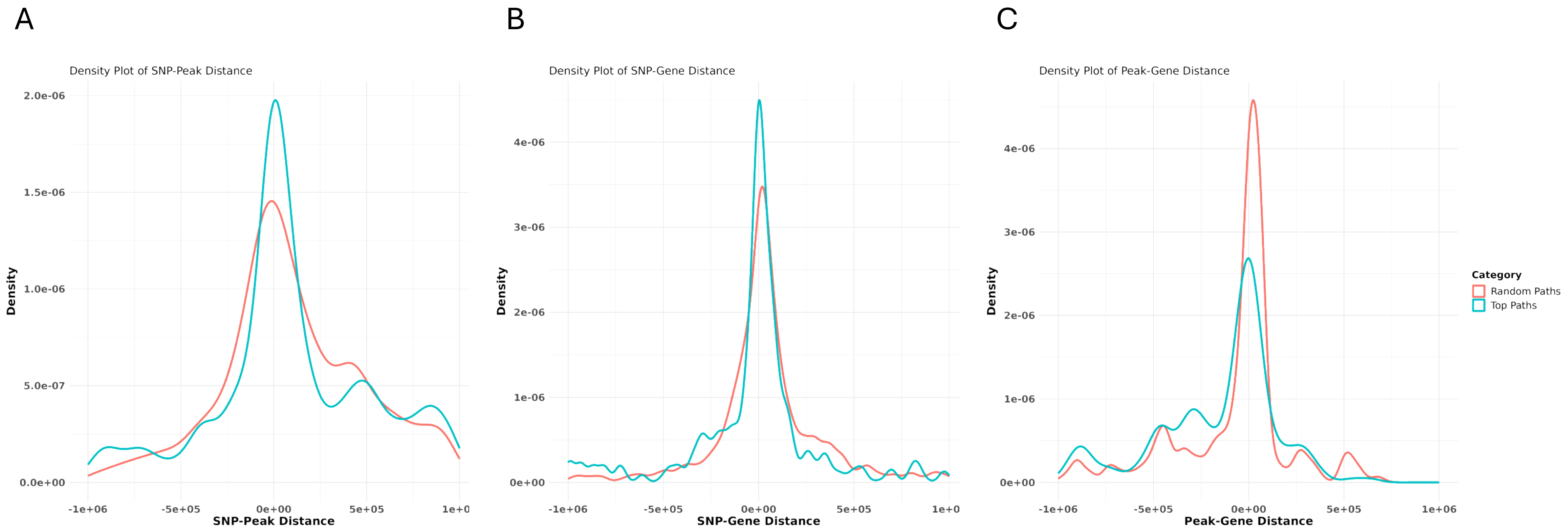
